# Epidermal expression of a sterol biosynthesis gene regulates root growth by a non-cell autonomous mechanism in *Arabidopsis*

**DOI:** 10.1101/204065

**Authors:** Eleri Short, Margaret Pullen, Gul Imriz, Dongbin Liu, Naomi Cope-Selby, Andrei Smertenko, Patrick J. Hussey, Jennifer F. Topping, Keith Lindsey

## Abstract

The epidermis has been hypothesized to play a signalling role during plant development. One class of mutants showing defects in signal transduction and radial patterning are those in sterol biosynthesis. The expectation is that sterol biosynthesis is a constitutive cell-autonomous process for the maintenance of basic cellular functions. The *HYDRA1* (*HYD1*) gene of Arabidopsis encodes an essential sterol Δ8-Δ7 isomerase, and although *hyd1* mutant seedlings are defective in radial patterning of several tissues, we show that the *HYD1* gene is expressed primarily in the root epidermis. Cell type-specific transgenic activation of *HYD1* transcription reveals that *HYD1* expression in the epidermis of *hyd1* null mutants is sufficient to rescue root patterning and growth. Unexpectedly, expression of *HYD1* in the vascular tissues and root meristem, though not endodermis or pericycle, also leads to phenotypic rescue. Phenotypic rescue is associated with rescued patterning of the PIN1 and PIN2 auxin efflux carriers. The importance of the epidermis is in part due to its role as a site for tissue-specific sterol biosynthesis, and auxin is a candidate for a non-cell autonomous signal.

## Introduction

A key question in plant development is how tissue patterning and cell expansion are coordinated in the course of organ growth. In the root of *Arabidopsis*, for example, the radial pattern is highly stereotyped, with predictable numbers of cells in each concentric layer of tissue, with coordination of cell expansion in each layer as the root grows (Dolan et al., 1993). This coordination of cell number and expansion is necessary, since plant cells are immobile and attached to each other. A failure of the coordination would likely to lead to growth and patterning defects.

The nature of such a coordination remains poorly understood. Mutant screens have led to the identification of genes essential for correct radial pattern in the *Arabidopsis* root, providing some insight into the molecular mechanisms involved. For example, the SCARECROW (SCR)/SHORTROOT (SHR) module controls ground tissue formation, and is characterized by the movement of the SHR protein from one cell layer (the stele), in which the gene is transcribed, to the cortex, where it regulates cell identity (Nakajima et al., 2001). The *keule* and *knolle* mutants exhibit radial defects such as bloated epidermal cells and very short roots, though the genes, that encode interacting components of the membrane trafficking system required for cytokinesis and cell wall construction, are expressed in all cells (Waizenegger et al., 2000; Assaad et al., 2001). Laser ablation experiments also highlight the importance of positional information in the regulation of root tissue patterning through as yet poorly defined signalling mechanisms (van den Berg et al., 1995, 1997). Non-autonomous signalling processes in radial patterning include the movement of transcription factors such as SHR between layers, and brassinosteroid (BR) signalling from the shoot epidermis has been implicated in regulating development of the ground and vascular tissues (Savaldi-Goldstein et al., 2007), and leaf shape (Reinhardt et al, 2007). The CRINKLY4 (CR4) receptor kinase of maize is expressed in the leaf epidermis and appears to signal to mesophyll cells, probably through an indirect mechanism (Jin et al., 2000; Becraft et al., 2001). In the root, epidermis-derived BR signalling also controls meristem size, through the modulation of, for example, the expression of the MADS-box transcription factor AGL42 in the quiescent centre (Hacham et al., 2011), though the transmitted signal remains unknown.

One class of mutants that exhibit radial patterning defects include the *hydra1*, *fackel/hydra*2 and *sterol methyltransferase* (*smt*) mutants, defective in sterol biosynthesis (Topping et al., 1997; Jang et al., 2000; Schrick et al., 2002, 2004; Souter et al., 2002). These are distinct from BR mutants, in that they cannot be rescued by exogenous supply of BRs (Topping et al., 1997; Schrick et al., 2000). Sterols are required for controlling membrane fluidity and permeability, and influence the activities of membrane-bound proteins (Grandmougin-Ferjani et al., 1997; Hartmann, 1998). Sterols are also implicated in the correct trafficking and localization of transporter proteins such as the PIN auxin efflux carriers (Willemsen et al., 2003; Men et al., 2008; Pan et al., 2009; Pullen et al., 2010), and in cell plate construction (Peng et al., 2002; Schrick et al., 2004). Given the water-insolubility and presumed lack of mobility of sterols for thermodynamic reasons, the expectation is that they are synthesised in all or the majority of cells, permitting basic cellular functions; and would function in a cell-autonomous fashion. There is also a proposed role for HYD1 in miRNA function, with a requirement of ARGONAUTE1 (AGO1) activity being dependent on sterol-dependent membrane composition (Brodersen et al., 2012). Mutants such as *hyd1*, *fk/hyd2* and *smt1* exhibit significant patterning and growth defects in most cell types, and are typically seedling-lethal, suggesting essential roles in all cell types (Jang et al., 2000; Schrick et al., 2002; Souter et al., 2002; Willemsen et al., 2003).

To investigate cell autonomy of sterol action in *Arabidopsis*, we investigated *HYD1* (At1g20050) expression and used transgenic activation systems to drive expression in different root cell types in the mutant background, to determine the cell types in which its expression is required for correct root development.

## Results and Discussion

*hyd1* seedlings exhibit abnormal morphogenesis and cell patterning and growth in the root (Fig. 1; Topping et al., 1997; Souter et al., 2002). The first evidence of defective radial pattern is seen during embryogenesis (Topping et al., 1997; Souter et al., 2002, 2004). Mutant seedlings typically develop multiple cotyledons of aberrant shape, a short hypocotyl and short root (Fig. 1B). The root has a defective apical meristem associated with aberrant patterning of surrounding cells (epidermis, columella, ground tissue, vascular tissue; Fig. 1C,D). Given the evident defects across several cell types, and the expectation that all cells contain sterols, we monitored spatial activity of the expression of a 2 kb fragment of the *HYD1* gene promoter as a transcriptional fusion reporter with a β-glucuronidase (GUS, *uidA*) gene in transgenics. The expression pattern in the primary root of transgenics is shown in Fig. 1E. Unexpectedly, results show a spatially restricted expression pattern, localized to the lateral root cap, epidermis of root elongation zone and, less strongly, in the root differentiation zone (especially in trichoblast files), with some detectable expression in the root cortex. This is consistent with cell expression profiling visualised in the Toronto expression profiling browser tool (http://bar.utoronto.ca/eplant/; Winter et al., 2007) based on data from Brady et al., (2007), (Supplementary Fig. 1). These observations raise the question of how such a localized expression pattern leads to radial patterning defects across a wider range of tissues in the primary root of the *hyd1* mutant.

**Fig. 1.**
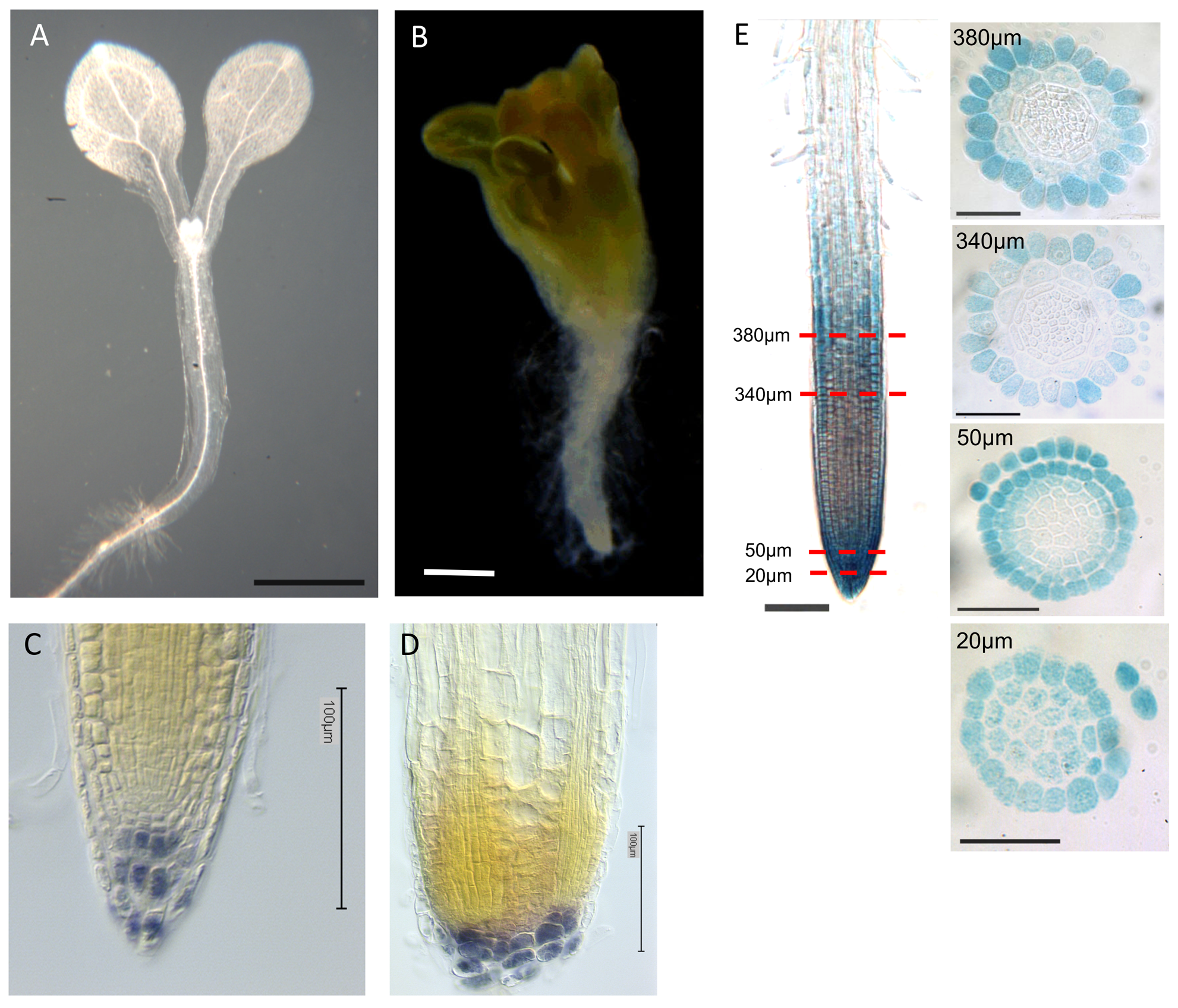
The *hyd1* mutant and *HYD1* expression. A. Wild type seedling, 6 dpg. Bar = 1 mm. B. *hyd1* mutant seedling, 6 dpg. Bar = 1 mm. C. Wild type seedling root stained with lugol, 6 dpg. Bar = 100 μm. D. *hyd1* seedling root stained with lugol, 6 dpg. Bar = 100 μm. E. proHYD1::GUS expression in wildtype seedling root at 7 dpg. Measurements in μm indicate the section distance from the root apex. Bar = 50 μm.

While sterols are transported between intracellular membrane compartments via lipid transfer proteins (Saravanan et al., 2009), there is no evidence that they are transported between cells. The question then is, how can localized expression of the *HYD1* gene, which is essential for radial patterning throughout embryogenesis and post-embryonic growth, mediate the development of cell types in which it is not active? We have shown, for example, that vascular patterning is abnormal, and PIN1 localization is aberrant in the vascular cells, even though *HYD1* is not expressed in those cells (Pullen et al., 2010). This suggests that a non-cell-autonomous signal is transferred from *HYD1*-expressing cells to non-expressing cells, to mediate wild-type tissue patterning.

To understand which cells might be sufficient and/or necessary for *HYD1* expression to mediate processes essential for root growth and development, the full length *HYD1* coding sequence (Topping et al., 1997) was cloned behind a variety of promoters and the UAS for use in mGAL4-VP16-GFP enhancer trap transactivation system (Laplaze et al., 2005), to drive *HYD1* transcription in different root cell types: the columella and QC, epidermis, endodermis, pericycle and vascular cells. Promoter::HYD1 and UAS::HYD1 fusions were transformed into wild-type *Arabidopsis*, crossed with the *hyd1* heterozygotes, and prospective homozygous mutant seedlings containing the promoter/UAS::HYD1 fusions were identified by genotyping and microscopy, for further analysis. Several independent transgenics were generated and typical expression patterns were identified in specific lines. For expression in the columella and QC we used the *POLARIS* (*PLS*) promoter (Casson et al., 2002) and the synthetic promoter DR5 (Sabatini et al., 1999), and the respective promoter-GUS expression patterns in both wild-type and *hyd1* mutant root tips are shown in Fig. 2A-D. For epidermal and lateral root cap expression we used the GAL4 driver line J2551, shown in wild-type and *hyd1* mutant roots in Fig. 2E-G; for endodermis, line J3611 (Fig. 2H-J); for pericycle line J0272 (Fig. 2K-M); and for vascular cells, line J0661 (Fig. 2N-P).

**Fig. 2.**
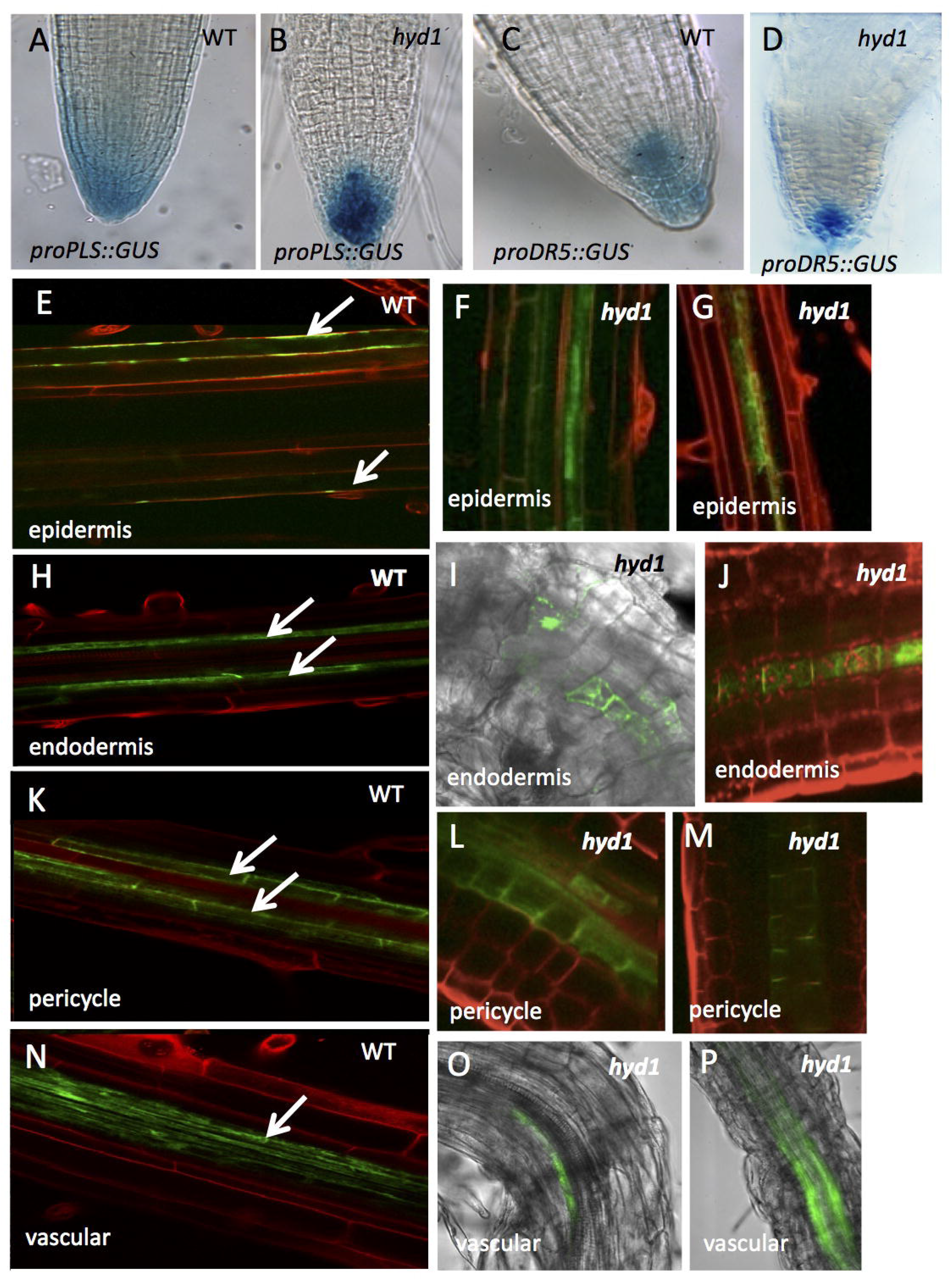
Cell type expression of promoters in wild type and *hyd1* mutants. A, B. proPLS::GUS expression in roots of wildtype (A) and *hyd1* (B) seedling primary root tips at 7 dpg. C,D. DR5::GUS expression in roots of wildtype (C) and *hyd1* (D) seedling primary root tips at 7 dpg. E-G. GFP expression in epidermal cells in GAL4 driver line J2551 in wildtype (E) and *hyd1* (F,G) seedling roots at 7 dpg. H-J. GFP expression in endodermal cells in GAL4 driver line J3611 in wildtype (H) and *hyd1* (I,J) seedling roots at 7 dpg. K-M. GFP expression in pericycle cells in GAL4 driver line J0272 in wildtype (K) and *hyd1* (L,M) seedling roots at 7 dpg. N-P. GFP expression in vascular cells in GAL4 driver line J0661 in wildtype (N) and *hyd1* (O,P) seedling roots at 7 dpg.

Expression in the *hyd1* mutant reflects the aberrant tissue patterning, but both promoter::GUS and UAS::HYD1:GFP lines were identified that exhibited expression in the expected cell types. Examples are given in Fig. 2 of different seedling lines exhibiting the various UAS::HYD1:GFP expression patterns. These results provide the basis for the use of the promoter/UAS regulatory sequences to drive *HYD1* transcription in specific cell types, to determine the effects on root development.

Genetically homozygous mutant seedlings expressing the *HYD1* coding region in different cell types were grown for up to 21 days on vertical agar plates for phenotypic and growth analysis. While the mean length of wild-type roots was ca. 12.4 cm at 14 dpg, and for the *hyd1* homozygous mutants was typically ca. less than 0.5 cm at 14 dpg, there were significant differences in the effects of expressing *HYD1* in different cell types (Fig. 3A,B). The most significant restoration of primary root growth was in *hyd1* seedlings expressing the *HYD1* gene in the epidermis (i.e. J2551>>HYD1; mean primary root length 9.1 cm at d 14), in the vascular tissues (i.e. J0661>>HYD1; mean primary root length 3.2 cm at d 14) and in the root columella and lateral root cap (DR5::HYD1, PLS::HYD1; mean primary root length up to ca. 7.8 cm at d 14). Therefore in each of these lines, primary root length was restored to ca. 60-80% that of wild-type by 14 dpg for epidermal expression, and with only ca. 30% wildtype growth seen with *HYD1* expression in vascular tissue (Fig. 3A,C). 55-65% of wild-type root growth was seen in seedlings with expression in the root columella and lateral root cap (Fig. 3B,D,E). Associated with significant restoration of root morphology and growth is an improved patterning of cells in the root tip, seen as a regularized organization of the starch-containing columella, though still with variable tiers of cells (Fig. 3F).

**Fig. 3.**
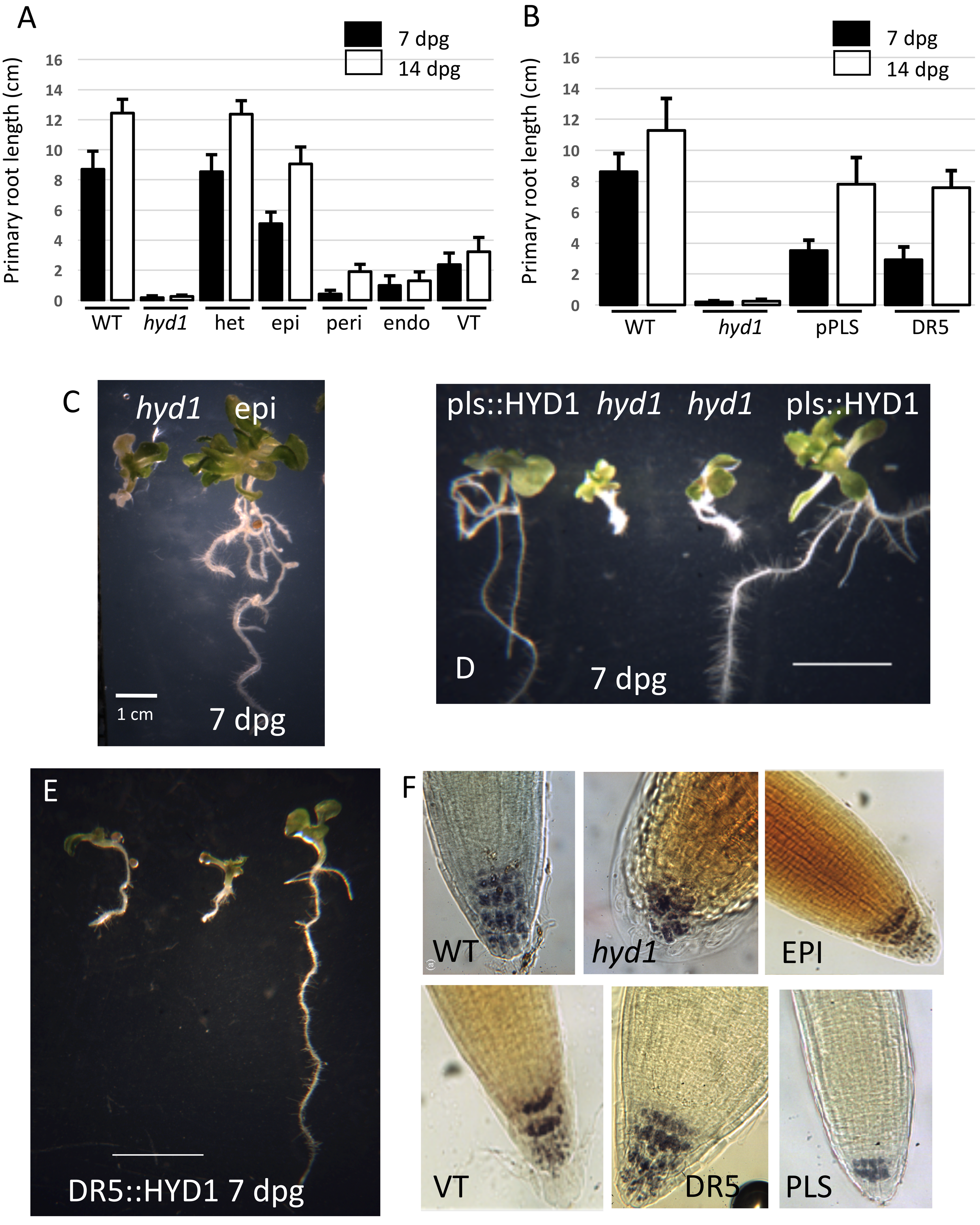
Effect of cell-type specific expression of the *HYD1* gene on root growth and columella organization. A. Primary root length of wildtype (WT), *hyd1*/WT heterozygous (HET), *hyd1* homozygous (*hyd1*) and transgenic *hyd1* seedlings expressing *HYD1* in epidermis (epi), pericycle (peri), endodermis (endo) and vascular tissues (VT) at 7 and 14 dpg. B. Primary root length of wildtype (WT), *hyd1* homozygous (*hyd1*) and transgenic *hyd1* seedlings expressing *HYD1* in root tips under the control of the PLS and DR5 promoters at 7 and 14 dpg. C. *hyd1* mutant and transgenic *hyd1* seedlings expressing *HYD1* in the GAL4 driver line J2551 in epidermal cells at 7 dpg. Bar = 1 cm. D. *hyd1* mutant and transgenic *hyd1* seedlings expressing proPLS::HYD1 in root tips at 7 dpg. Bar = 1 cm. E. *hyd1* mutant and transgenic *hyd1* seedlings expressing DR5::HYD1 in root tips at 7 dpg. Bar = 1 cm. F. Root tips of wildtype (WT), *hyd1* and transgenic *hyd1* seedlings expressing *HYD1* in the GAL4 driver line J2551 in epidermis (EPI), in line J0661 in vascular tissues (VT) and in root tips under the control of the DR5 and PLS promoters at 7 dpg.

Root growth and cell patterning, including columella organization, has been linked to auxin concentration and response in the root (Sabatini et al., 1999; Aida et al., 2004). Sterols have been shown to be required for correct auxin-mediated gene expression (Souter et al., 2002, 2004) and for PIN localization, including in the *hyd1* and *smt* mutants (Carland et al., 2010; Pullen et al., 2010). PIN-FORMED (PIN) proteins act as auxin efflux carriers, allowing auxin to be transported in a directional manner to establish gradients across tissues, often with developmental or tropic (e.g. gravitropic) consequences. Correct membrane sterol composition has been shown to be required for correct PIN2 polarity and gravitropic response (Men et al., 2008). To determine whether the activation of the *HYD1* gene in specific cell types was associated with a restoration of PIN localization, PINs 1 and 2 were immunolocalized in both the mutant, wild-type, and transgenic lines expressing *HYD1* under control of cell type-specific promoters.

Results presented in Fig. 4 show that, as expected, the wild-type root tip shows PIN1 localized to the basal region of cells in the stele of the root, and PIN2 was localized to the apical side of epidermal cells (Fig. 4A). In the *hyd1* mutant, both abnormal cellular patterning and loss of polar PIN1 and 2 localization are evident (Fig. 4B). Cellular patterning in seedlings expressing *HYD1* in the vascular tissues (J0661>>HYD1) (Fig. 4C) or pericycle (J0272>>HYD1) (Fig. 4D) is poorly restored, and PIN expression and localization is variable, associated with relatively poor primary root growth (Fig. 3A). In seedlings expressing *HYD1* in the epidermis (J2551>>HYD1), cellular patterning is similar to wild-type, as is the localization of PIN2 and PIN1 (Fig. 4E). This is associated with relatively long primary roots in these seedlings (Fig. 3A). In proPLS::HYD1 seedlings (Fig. 4F), radial patterning of the root is restored close to wild-type, with an improvement of PIN localization compared to either the *hyd1* mutant or, for example, the vascular tissue line (J0661>>HYD1). The expression of two auxin-regulated genes, *IAA1* and *IAA2*, are known to be poorly expressed in the *hyd* mutants (Souter et al., 2004) and activation on *HYD1* in the epidermis (J2551>>HYD1), and also to some extent in the vascular tissues (J0661>>HYD1) and root tip (PLS::HYD1) leads to some recovery of the expression levels of both genes (Fig. 4G).

**Fig. 4.**
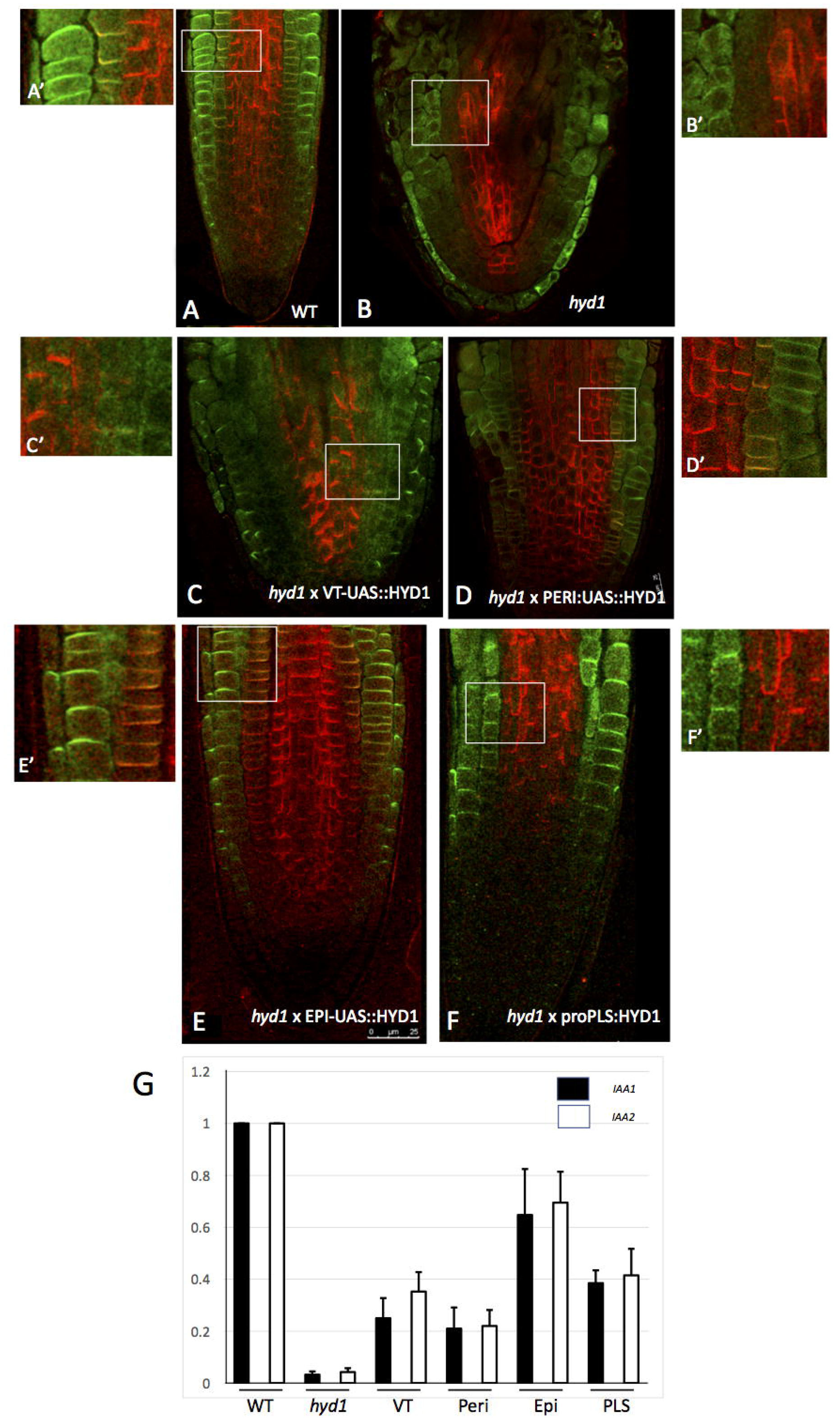
Effect of cell-type specific expression of the *HYD1* gene on PIN proteins and auxin gene expression. PIN1 (red) and PIN2 (green) immunolocalization in wildtype root (A, A'), in *hyd1* mutant root (B, B'), in *hyd1* mutant root expressing *HYD1* in the GAL4 driver line J0661 in vascular tissues (VT) (C, C'), in *hyd1* mutant root expressing *HYD1* in the GAL4 driver line J0272 in pericycle cells (D, D') in *hyd1* mutant root expressing *HYD1* in the GAL4 driver line J2551 in epidermal cells (E, E') and in *hyd1* mutant root expressing *HYD1* under the control of the PLS gene promoter (F, F'), at 7 dpg. G. Expression of *IAA1* (black bars) and *IAA2* (open bars) in wildtype (WT), *hyd1* mutant (HYD1), vascular tissues (VT), pericycle cells (Peri), epidermal cells (Epi) and root tip (PLS) relative to wildtype (value 1), determined by qRT-PCR. Means α SD of 4 biological replicates.

The evidence presented here suggests that the epidermis plays an important role in controlling growth through its role as a site for sterol biosynthesis, and this involves a non-autonomous signalling pathway, for which auxin is a strong candidate. Previous evidence demonstrated a non-autonomous role for the epidermis in BR synthesis, but the signal involved was not identified. Our data suggest that at least one coordinating signal across tissues is auxin, and the role of sterols in this context is to mediate correct localization and function of the PIN proteins, which are responsible for directional auxin transport. Sterols are known to control PIN polarization, and the data presented show that sterols are likely functioning in a non-autonomous fashion by mediating auxin gradient establishment, which in turn controls patterning and growth, through e.g. activation of the PLT/WOX5 mechanism (Aida et al., 2004; Sarkar et al., 2007). Exogenous auxin does not rescue the *hyd* mutant phenotype, supporting the view that gradients of auxin rather than absolute levels are required for correct development; and the *hyd1* mutant lacks PIN3 proteins (Souter et al., 2002), which in wild-type accumulate distal to the quiescent centre and distribute auxin both down into the columella and laterally towards the epidermis and cortex (Friml et al., 2002). *hyd1* also shows defective PIN1 and PIN2 localization and auxin patterning, associated with defective cell patterning (Pullen et al., 2010). The epidermis appears critical as a site of sterol biosynthesis via HYD1 - the *hyd1* mutants fail to accumulate key sterols (Souter et al., 2002), and expression of the *HYD1* gene specifically in the epidermis significantly rescues root growth and patterning of cells in the root tip.

Interestingly, *HYD1* expression in the root cap (both columella and lateral root cap cells) also leads to significant root growth rescue, presumably by promoting PIN activity there to ensure stem cell niche activity - columella patterning is rescued, as well as root growth. Expression in the pericycle, endodermis or vascular tissues, on the other hand, has limited effects on root growth (Fig. 3), pointing to a mechanism distinct to, for example, the role of gibberellins in the endodermis (Ubeda-Tomas et al., 2008, 2009). These results show that the role of the epidermis in regulating root growth can at least in part be explained by its role in non-autonomous auxin signalling via sterol biosynthesis.

## Materials and Methods

### Plant material

The *hyd1* mutant was identified in an insertional mutagenesis screen as described previously (Topping et al., 1997; Souter et al., 2002). The full length *HYD1* cDNA sequence was cloned into the vector pCIRCE, and fused to the promoters DR5 (Sabatini et al., 1999), PLS (Casson et al., 2002) or UAS (Laplaze et al., 2005). The constructs were introduced into *Arabidopsis thaliana* by floral dip transformation (Clough and Bent, 1998). Homozygous T2 lines containing the UAS:HYD1 were crossed with the GAL4 driver lines J2551 (epidermal and lateral root cap expression); J3611 (endodermis); J0272 (pericycle); and J0661 (vascular cells), kindly provided by Dr. Jim Haseloff (Cambridge University, UK). Plants homozygous for all HYD1 constructs were crossed respectively with plants heterozygous for the *hyd1* mutation, and selfed to identify progeny that was homozygous for both the original *hyd1* mutation and the *HYD1* fusion transgene, for further analysis. For growth assays, seeds were stratified, surface sterilized and grown on vertical agar plates containing half-strength Murashige and Skoog medium as described previously (Topping et al., 1997).

### Gene expression analysis

RNA was extracted from seedlings, and gene expression measured by quantitative RT-PCR, with *ACTIN3* as an internal standard, as described previously (Rowe et al., 2016).

### Immunofluorescence microscopy and imaging

*Arabidopsis* roots were fixed for 60 min at room temperature with 4% (w/v) paraformaldehyde in 0.1 M Pipes, pH 6.8, 5 mM EGTA, 2 mM MgCl_2_, and 0.4% Triton X-100. The fixative was washed away with PBST buffer, and cells were treated for 8 min at room temperature with the solution of 2% (w/v) Driselase (Sigma) in 0.4 M mannitol, 5 mM EGTA, 15 mM MES, pH 5.0, 1 mM PMSF, 10 μg mL^−1^ leupeptin and 10 μg mL^−1^ pepstatin A. Thereafter roots were washed two times 10 min each in PBST and in 1% (w/v) BSA in PBST for 30 min, and incubated overnight with a primary antibody. The primary antibodies rabbit anti-PIN1 (1:150) and guinea pig anti-PIN2 (1:150). Specimen were then washed three times for 90 min in PBST and incubated overnight with goat anti-mouse TRITC and anti-rabbit FITC conjugated secondary antibodies diluted 1:200. After washing in the PBST buffer, specimens were mounted in the Vectashield (Vector Laboratories, Burlingame, CA) mounting medium. Images were acquired using Leica SP5 Laser Confocal Scanning Microscope using excitation at 488 nm line of Argon laser for FITC or 561 nm excitation of solid-state laser for TRITC. The emitted light was collected at 505-550 nm or 570-620 nm respectively.

Light micrographs were acquired using a Zeiss Axioskop microscope (Carl Zeiss Ltd, Herts, UK) equipped with Photometrics COOLSNAP^TM^cf colour digital camera (Roper Scientific Inc., Trenton, New Jersey, USA) and OpenLab3.1.1 software (Improvision, Coventry, UK). GFP signal in roots was imaged using Leica SP5 confocal microscope using 488 nm line of argon laser and emission was collected between 505 and 530 nm. The roots were mounted in double distilled water under a large (25×50 mm) zero-thickness coverslip.

## Acknowledgements

The authors are grateful to Prof. Klaus Palme (University of Freiburg) for providing antibodies against PIN1 and PIN2.

## Author contributions

KL and JT conceived the project, ES, MP, GU, DL, N C-S, AS carried out experimental work, KL wrote the manuscript, JT, AS and PJH revised the manuscript.

---

**Supplementary Fig. 1.** *HYDRA1* gene expression visualised in the Toronto expression profiling browser tool (http://bar.utoronto.ca/eplant/; Winter et al., 2007) based on data from Brady et al. (2007). Expression is shown in untreated roots, and in roots subjected to salt stress, iron deficiency and nitrogen application.

